# DnaA modulates the gene expression and morphology of the Lyme disease spirochete

**DOI:** 10.1101/2024.06.08.598065

**Authors:** Andrew C. Krusenstjerna, Nerina Jusufovic, Timothy C. Saylor, Brian Stevenson

**Affiliations:** Department of Microbiology, Immunology, and Molecular Genetics, University of Kentucky College of Medicine, Lexington, Kentucky, USA; Department of Entomology, University of Kentucky, Lexington, Kentucky, USA

## Abstract

All bacteria encode a multifunctional DNA-binding protein, DnaA, which initiates chromosomal replication. Despite having the most complex, segmented bacterial genome, little is known about *Borrelia burgdorferi* DnaA and its role in maintaining the spirochete’s physiology. In this work we utilized inducible CRISPR-interference and overexpression to modulate cellular levels of DnaA to better understand this essential protein. Dysregulation of DnaA, either up or down, increased or decreased cell lengths, respectively, while also significantly slowing replication rates. Using fluorescent microscopy, we found the DnaA CRISPRi mutants had increased numbers of chromosomes with irregular spacing patterns. DnaA-depleted spirochetes also exhibited a significant defect in helical morphology. RNA-seq of the conditional mutants showed significant changes in the levels of transcripts involved with flagellar synthesis, elongation, cell division, virulence, and other functions. These findings demonstrate that the DnaA plays a commanding role in maintaining borrelial growth dynamics and protein expression, which are essential for the survival of the Lyme disease spirochete.

**IMPORTANCE:** Lyme disease is the most prevalent tick-borne infection in the Northern Hemisphere. *Borrelia burgdorferi*, the causative spirochete bacteria, has been maintained in nature for millennia in a consistent enzootic cycle between *Ixodes* ticks and various small vertebrate hosts. During the tick’s blood meal, *B. burgdorferi* substantially increases its replication rate, alters its repertoire of outer surface proteins, and disseminates into the new vertebrate host. Across eubacteria, DnaA is the master regulatory protein that initiates chromosomal replication and acts as a transcription factor to regulate specific pathways. Here, we describe the roles that *B. burgdorferi* DnaA has on the physiology and gene expression of this medically important pathogen.

## Introduction

Replication plays a significant role in the enzootic lifecycle of *Borrelia burgdorferi*, the Lyme disease spirochete. An extracellular bacterial parasite, *B. burgdorferi* exclusively colonizes Ixodid ticks and assorted vertebrates (1, 2). Naïve tick larvae acquire the Lyme bacteria when they feed on an infected host. After this single blood meal, the larvae drop off, overwinter, and molt to their nymphal stage. During this time, *B. burgdorferi* persists within the midgut of the nutrient-deplete tick. Once the nymph emerges and takes its blood meal, the spirochetes obtain the requisite materials to replicate, divide, and disseminate from the tick to the new vertebrate host (3, 4).

The highly conserved AAA+ family protein DnaA initiates the replication of bacterial chromosomes (5–7). DnaA monomers recognize the *oriC* locus and cooperatively multimerize to form a helical structure. The DnaA filament promotes the separation of AT-rich DNA elements, which recruits helicase and replication machinery. DnaA not only initiates chromosomal replication but also binds elsewhere in the genome to regulate gene expression (8, 9). We recently found that *B. burgdorferi* DnaA directly regulates the *dnaX-ebfC* operon, which codes for the Tau (*τ*) subunit of DNA polymerase III holoenzyme, DnaX, and a regulatory nucleoid-associated protein, EbfC (10). These genes are highly expressed during periods of rapid spirochete replication (10–12). At the tick-vertebrate interface, we hypothesize that *B. burgdorferi* DnaA not only commits to its replication initiation function but coordinates the expression of genes needed for vertebrate infection (10, 13, 14).

Replication of the *B. burgdorferi* linear chromosome proceeds bidirectionally from the centrally-located *oriC* (15). Outside of this, little is known about borrelial DNA replication, its regulation, or coordination with other cellular processes. Many of the genes involved in those pathways are essential, which has hampered their investigation. Historically, studying essential loci required the development or isolation of conditional mutants. The first studies into bacterial DnaA proteins utilized temperature-sensitive mutants (16). In the era of molecular genetics, many tools have been developed to produce conditional phenotypes more readily, such as inducible promoters. Overexpression has been a reliable means of elucidating a protein’s function in *B. burgdorferi* and other bacteria (14, 17, 18). Recently, inducible CRISPR interference (CRISPRi) systems have been developed and added to the repertoire of tools for studying borrelial biology (19, 20). This work will describe the consequence of CRISPRi-mediated knockdown of the essential replication initiator protein DnaA in *B. burgdorferi*. Using this approach, coupled with overexpression, we observed profound consequences of DnaA-dysregulation on spirochete morphology, chromosome partitioning, and gene expression.

## RESULTS

### Construction and validation of plasmids to dysregulate levels of DnaA in *B. burgdorferi*

DnaA is essential for initiating chromosomal replication, so deleting DnaA is lethal to bacteria. Thus, to gain insight into the impacts of DnaA on borrelial physiology, we produced IPTG-inducible constructs to reduce or elevate cellular levels of DnaA. Knockdown was carried out using the newly refined all-in-one CRISPR interference (CRISPRi) shuttle vector, which places the expression of both the sgRNA and a borrelial codon-optimized dCas9 gene under the control of the Lac repressor (20). We designed two CRISPRi constructs with sgRNAs targeting either the template (*dnaA*_T1_) or non-template (*dnaA*_NT1_) DNA strands directly 5’ or within the *dnaA* gene (**Fig. 1A**). Overexpression was achieved using plasmid pACK121, which contains an inducible *dnaA* with an N-terminal 3xFLAG tag. All constructs were transformed into *B. burgdorferi* strain B31-e2, hereafter referred to as e2. The spirochetes tolerated each construct well, replicating at rates comparable to the parental strain without the inducer (**Fig. 1B**), with one exception. The *dnaA*_T1_ CRISPRi strain replicated slightly slower than the others, although it reached the same final density. This may suggest leakiness of the hybrid *lac* promoter that controls the sgRNA and dCas9 gene (B. Murphy & W. Zückert, personal communication).

**Figure 1.**
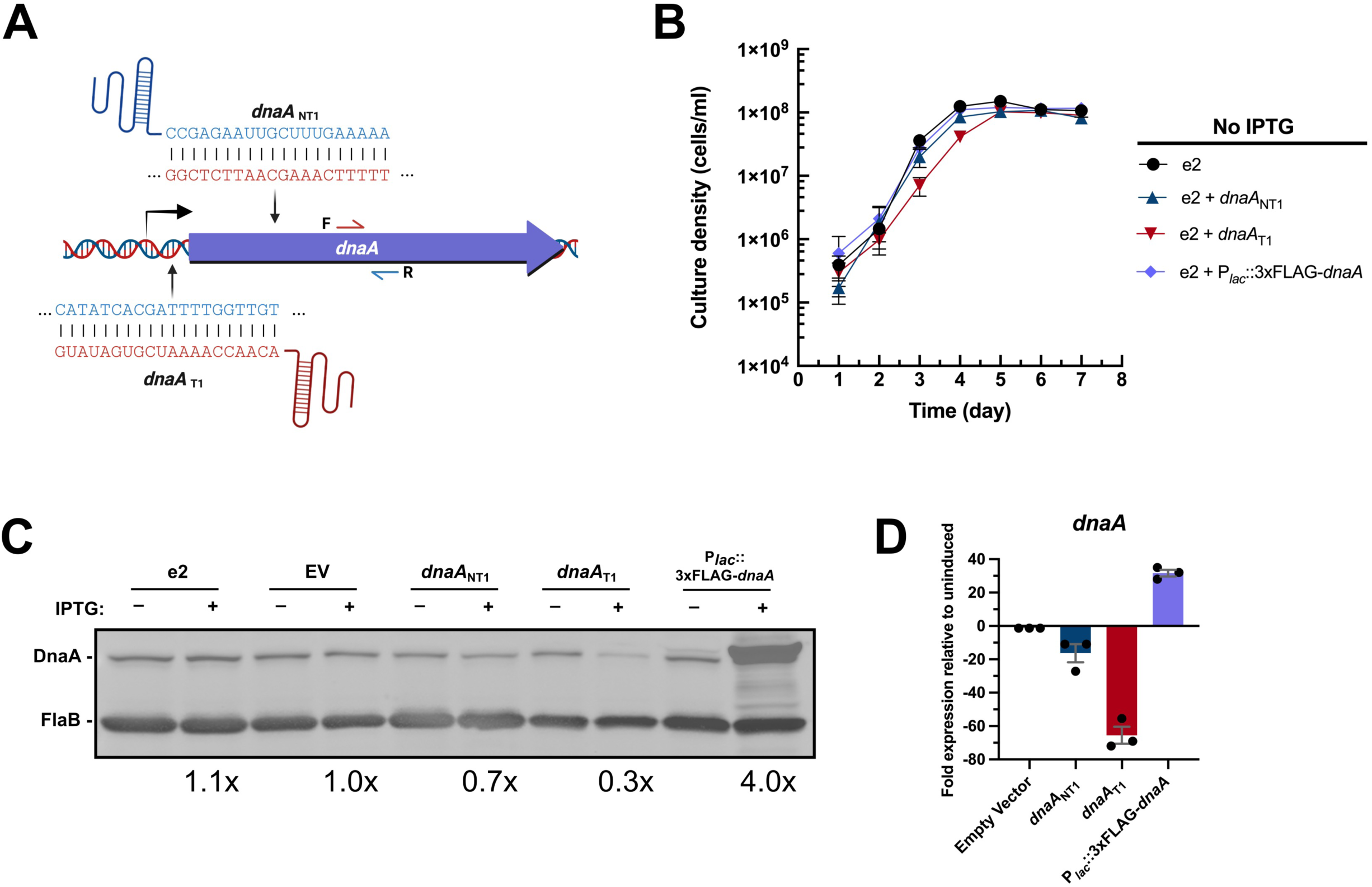
Validation of conditional dysregulation of DnaA. **(A)** Two sgRNAs were designed to knock down borrelial *dnaA* transcription by CRISPRi. One sgRNA targeted the template strand directly upstream of the ORF (*dnaA*_T1_), while the other targeted the template strand within the ORF (*dnaA*_NT1_). **(B)** The *dnaA* CRISPRi bacteria, CRISPRi empty vector bacteria, and the strain with the *dnaA* overexpression plasmid grew at the same rate as the parental e2. Adding IPTG to these strains resulted in appropriate knockdown or overexpression of DnaA protein **(C)** and transcript **(D)**, as assessed by immunoblot and qRT-PCR. The overexpression shuttle vector encodes a DnaA with an N-terminal 3xFLAG moiety and thus migrates above the native protein. Error bars in (B) and (D) represent the standard error of the mean (SEM).

To assess the efficacy of these inducible constructs, spirochetes of each strain were grown to mid-exponential growth phase (3-5ⅹ10^7^ cells/mL), then incubated overnight with 0.5 mM IPTG. The *dnaA*-targeting CRISPRi strains decreased the expression of DnaA, with the construct targeting the template strand, *dnaA*_T1_, consistently yielding the most effective gene silencing (**Fig. 1C** and **D**). The potency of the *dnaA*_T1_ construct likely explains the slowed spirochete growth seen in the absence of IPTG (**Fig. 1B**). Overexpression resulted in elevated quantities of DnaA protein and transcript (**Fig. 1C** and **D**). Neither the CRISPRi empty vector (EV) nor the parental e2 control strains yielded changes in DnaA levels when IPTG was added, demonstrating that we could specifically and effectively alter cellular DnaA concentrations.

### Proper DnaA expression is essential for borrelial growth

Given its essential nature for chromosomal replication, we first sought to evaluate the consequence of DnaA dysregulation on *B. burgdorferi* growth and cell division. To achieve this, all strains were grown to mid-exponential phase and passaged into fresh media with 0.5 mM of IPTG at a density of 1×10^5^ cells/mL. Bacterial numbers of each culture were counted every day for seven days to generate growth curves.

CRISPRi knockdown of DnaA levels substantially reduced the generation time and carrying capacity of *B. burgdorferi* (**Fig. 2A** and **B**). The severity of these phenotypes was consistent with the knockdown efficacy of the two constructs, with *dnaA*_NT1_ cultures reaching a maximum density that was one log lower and *dnaA*_T1_ cultures maxing out two logs lower than the controls. This suggests that reducing the cellular concentration of DnaA directly limits the number of division cycles, which is consistent with DnaA’s role as the chromosomal replication initiator.

**Figure 2.**
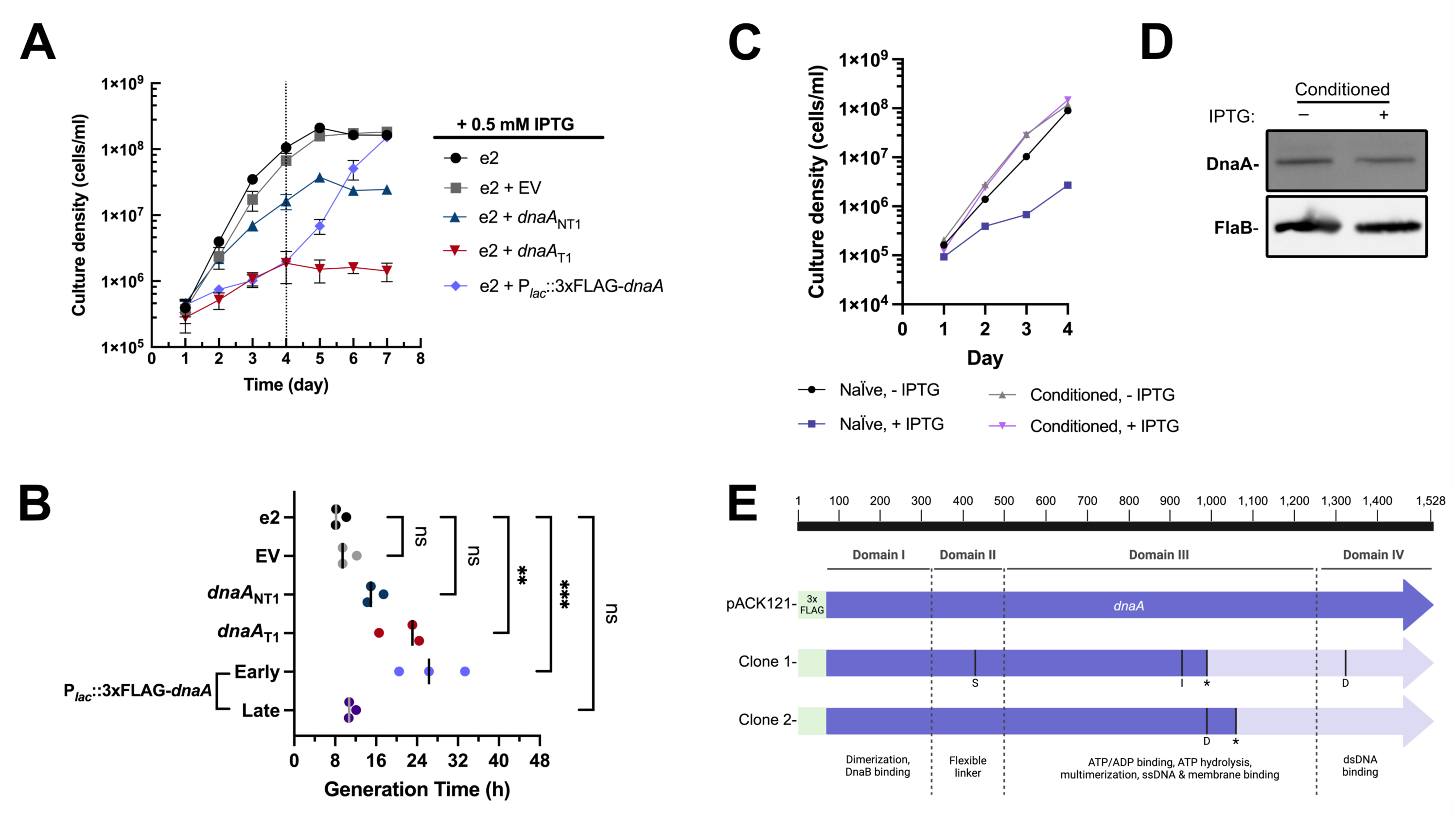
Appropriate expression of DnaA is required for *B. burgdorferi* replication. **(A)** Growth curve studies were conducted on different *B. burgdorferi* strains incubated with 0.5 mM IPTG. Three independent cultures of each strain were measured for growth curve analysis. Error bars represent the SEM. The dashed line indicates when the overexpression strain resumed growth. **(B)** The generation time was determined from the growth curve analyses from three individual cultures of each strain. For the overexpression strain, the growth rate was determined before (early; 1-4 dpi) and after the resumption of growth (late; 4-7 dpi). The resumption of replication in that strain was evidently due to mutation of the overexpressed *dnaA* gene. The p-values are indicated as follows: **, p ≤ 0.01; ***, p ≤ 0.001; ns, not significant, p > 0.05. **(C)** IPTG-treated DnaA overexpression spirochetes that resumed growth (conditioned) lost sensitivity to IPTG when passaged to fresh media and did not overproduce 3xFLAG-DnaA as assessed by immunoblot **(D**), supporting the conclusion that accumulated mutations in the overexpression *dnaA* permitted replication. **(E)** Sequenced plasmid from these conditioned spirochetes showed unique mutations (S = substitution; I= Insertion; D = deletion) in the 3xFLAG-*dnaA* ORF that caused a frameshift and truncation of the full-length protein. The location of the truncation is indicated by *.

Overproducing DnaA slowed growth during the first four days post-inoculation (dpi), growing at a rate similar to the *dnaA*_T1_ culture (**Fig. 2A** and **B**, early). After that time, however, cultures consistently resumed logarithmic growth, yielding generation times that were indistinguishable from the parental strain (**Fig. 2B**, late). We hypothesized this abrupt resumption of growth was due to mutations in the inducible *dnaA* plasmid that eliminated overexpression. To test this, we passaged spirochetes that had resumed growth into fresh media with IPTG to see if growth would still be perturbed. Consistent with our hypothesis, these “conditioned” bacteria immediately grew at rates that were similar to the control strain (**Fig. 2C**), and induction of the 3xFLAG tagged DnaA did not occur (**Fig. 2D**). To further test our hypothesis, we extracted the DnaA overexpression plasmid from the *B. burgdorferi* cultures and transformed them into *E. coli*. We sequenced purified plasmids from two colonies derived from the same induced borrelial culture. Each clone had unique mutations within the *dnaA* ORF that truncated the full-length protein (**Fig. 2E**). Notably, both mutations resulted in the complete loss of domain IV, the DNA-binding domain of DnaA. Plasmid isolated from transformed *B. burgdorferi,* which had never been exposed to IPTG, had no mutations within the inducible *dnaA*. These escape mutants, taken with the growth dynamics, highlight the stressful nature of *dnaA* overexpression, which has consistently been observed in other bacterial organisms (21–23).

Unlike the overexpression strain, we never observed a resumption of growth in the induced *dnaA* CRISPRi strains. Furthermore, normal growth was attainable from knocked-down cultures after passaging spirochetes into fresh media without IPTG. These collective data demonstrate that precisely controlled levels of DnaA are required for optimal borrelial growth.

### DnaA affects spirochete cell division/elongation

Chromosomal replication is intimately tied to cellular elongation and division (24–26). Having found that the conditional DnaA mutants had replication defects, we also sought to assess the impact of DnaA dysregulation on *B. burgdorferi* cell length. Over seven days, we imaged and measured spirochetes. In the parental e2 cultures, we observed a consistent pattern wherein median cell length peaked during early exponential phase, followed by a steady decrease, then stabilization once the cultures reached stationary phase (**Fig. 3A, 3E**, and **3I**).

**Figure 3.**
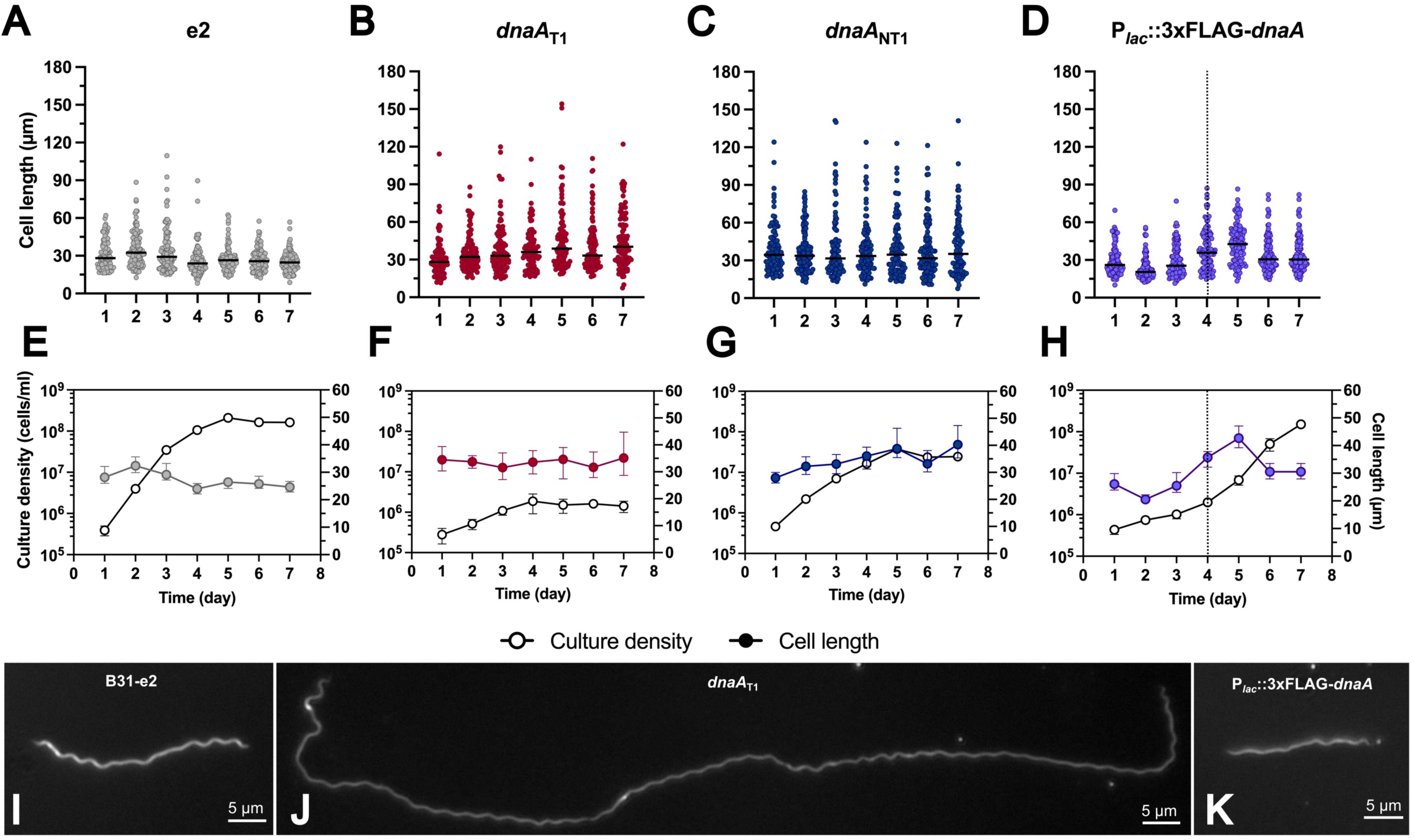
DnaA dysregulation affects spirochete cell length. **(A-D)** Cell lengths of the 0.5 mM IPTG-treated *B. burgdorferi* B31-e2 strains measured over seven days (gray = e2; red = *dnaA*_T1_; *dnaA*_NT1_ = blue; P*_lac_*::3xFLAG-*dnaA* = purple). The line represents the median cell length. **(E-H)** The median cell length (filled-in circle) with error bars representing the 95% CI superimposed against the corresponding growth curve (empty circle) from Fig. 2. The dashed line indicates when the overexpression strain resumed growth. **(I-K)** Representative images of observed phenotypes. **(I)** The e2 cell of approximate median length at 2 dpi. **(J)** DnaA-depleted cell with the characteristic elongation phenotype. **(K)** Median size of a DnaA overexpression spirochete at 2 dpi.

These findings sharply contrast with what we observed in the CRISPRi knockdown strains, where the spirochetes steadily maintained median lengths of 30-40 µm (**Fig. 3B-C, 3F, 3H**, and **3I**); the parental e2 strain had median lengths that spanned from 24-32 µm. The ranges (maximum-minimum) of the cell lengths within these knockdown strains (147 µm, *dnaA*_T1_; 134 µm, *dnaA*_NT1_) were greater than that of the overexpression (77 µm) and parental (101 µm) bacteria.

When DnaA was overexpressed, the median cell length decreased by 2 days post-induction (**Fig. 3D**, **3H**, and **3K**). As cultures accumulated mutations in the overexpressed *dnaA*, cell lengths gradually increased to a peak at day 5, then decreased. This oscillation in spirochete length aligned with the observed growth pattern of this strain, where slowed replication corresponded to cells of shorter size and rapid replication to longer. This suggests that high levels of DnaA can diminish borrelial cell length. These results demonstrate that DnaA plays a role in regulating *B. burgdorferi* cell elongation and/or division and that maintaining proper DnaA levels is critical to these processes.

### Impact of DnaA on ploidy and partitioning

*B. burgdorferi* is polyploid during exponential growth *in vitro* (27). As DnaA is the master initiator of chromosomal replication and affects *B. burgdorferi* morphology, we hypothesized that the ploidy and partitioning of chromosomes could be altered when DnaA levels are dysregulated. To test this, we transformed the parental e2 and *dnaA*_T1_ strains with a construct expressing the chromosomal ParB protein fused to mCherry (pBSV2G_P_0826_-mCherry*_Bb_*-ParB), which has been demonstrated to bind near *oriC* (27). We first grew the two strains to mid-exponential phase without IPTG and measured the length and ParB-*oriC* puncta per cell. The parental bacteria were smaller than those of the uninduced *dnaA*_T1_, consistent with our prior observations. Yet, both strains had similar numbers of ParB-*oriC* (**Fig. 4**). After taking these measurements, the *dnaA*_T1_ culture was split in half, and IPTG was added to one of the cultures. The parental, uninduced, and induced strains were then incubated and examined for three more days.

**Figure 4.**
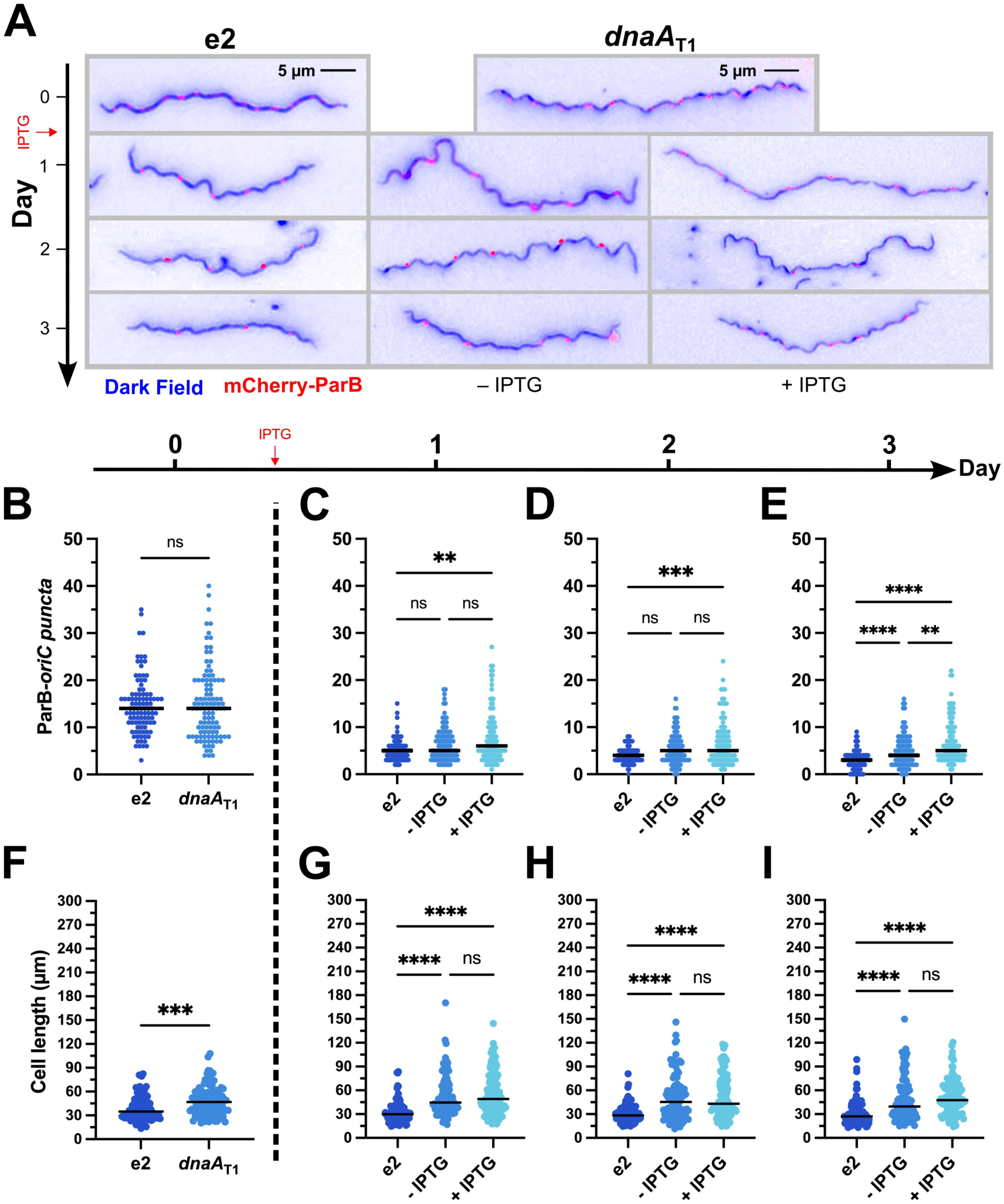
Knockdown of DnaA increases the number of chromosomes per spirochete. The *B. burgdorferi* CRISPRi *dnaA*_T1_ and parent B31-e2 strains were transformed with a construct encoding a mCherry-tagged ParB protein, which binds near the *oriC*, to identify the locations of the chromosomes. The two strains were grown without inducer until mid-log (*t* = 0), and then the *dnaA*_T1_ strain was divided into two cultures, and one of those was induced with 0.5 mM IPTG. Cultures were tracked for three more days. **(A)** Representative merged micrographs (blue = dark field; red = mCherry-ParB) of spirochetes with the approximate median numbers of ParB-*oriC* puncta **(B)** and cell length **(C)**. Significant differences in numbers of puncta and cell length at day 0 were determined by the Mann-Whitney U test. Differences between the three groups (e2, - IPTG, and + IPTG) on days 1-3 were tested for by the Kruskal-Wallis test and then Dunn’s test for multiple comparisons. Multiplicity-adjusted p-values are reported. **, p ≤ 0.01; ***, p ≤ 0.001; ****, p ≤ 0.0001; ns, not significant, p > 0.05. Approximately 50 spirochetes were imaged for each strain on each day. Data were gathered from two independent experiments.

On the first day post-induction, as the cultures entered stationary phase (≥1×10^8^ cells/mL), all strains experienced a dramatic decline in the total ParB-*oriC* puncta per cell, consistent with prior observations (27). Over the next two days, the number of these continued to decrease in the parental and uninduced cultures, whereas the induced culture plateaued and ended with significantly more ParB-*oriC* puncta. Furthermore, the induced cell lengths, while longer than the parental, were the same as the uninduced cells, indicating the difference in puncta wasn’t simply due to differences in cell length. With this, we concluded that depleting DnaA impairs the completion of borrelial chromosomal replication and cell division.

In addition to differences in the number of ParB-*oriC* puncta in the *dnaA*-conditional mutant, we also noted an apparent difference in their spacing (**Fig. 5A-B**) relative to the parental strain (**Fig. 5C**). Some cells had wide gaps between the foci, while others had foci that were close together. We assessed the cultures two- and three-days post-induction and found about 23.7% of parental e2 cells had irregular puncta spacing. The *dnaA*_T1_ CRISPRi strains, in contrast, had significantly greater proportions of 53.8% and 55.3% of spirochetes with irregular spacing without and with added IPTG, respectively (**Fig. 5D**). Thus, knocking down DnaA also perturbs chromosomal partitioning.

**Figure 5.**
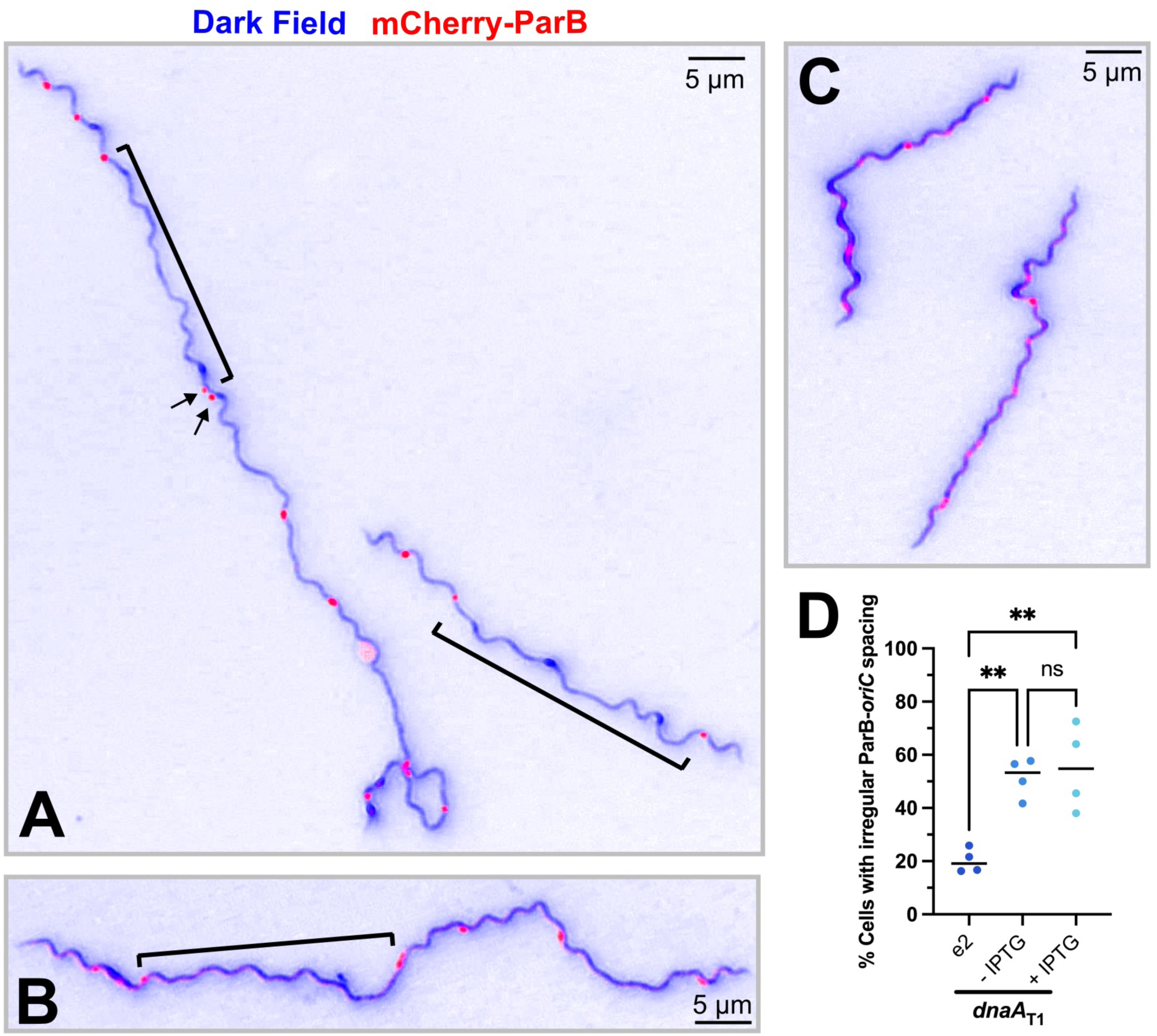
Irregular spacing of *oriC* in *dnaA* knockdown *B. burgdorferi*. **(A-B)** Representative images of spirochetes containing the *dnaA*_T1_ CRISPRi plasmid with irregular spacing of ParB-*oriC* puncta (red). DnaA-depleted cells had large regions where no foci were observed (brackets). Some cells also had foci that were close together (arrows). **(C)** Parental B31-e2 spirochetes with approximately the same number of ParB-*oriC* puncta as the mutant but exhibit regular oriC spacing. **(D)** The number of spirochetes with irregular ParB-*oriC* was quantified (e2, *n* = 184; uninduced *dnaA*_T1_, *n* = 198 cells; induced *dnaA*_T1_ *n* = 206). One-way ANOVA with Tukey’s post-hoc test was done to determine the significance and make comparisons. The multiplicity-adjusted p-values are reported **, *p* ≤ 0.01; ns, not significant, *p* > 0.05.

### Dysregulation of DnaA disrupts flagellar homeostasis

Depletion of DnaA also resulted in *B. burgdorferi* cells with aberrant helicity. Specifically, we observed spirochetes with a complete or partial loss of their characteristic corkscrew shape (**Fig. 6A** and **B**, respectively). Some of these tube-shaped cells were substantially elongated and showed evidence of incomplete division (**Fig. 6C** and **Supplemental Video**). About 13% of the spirochetes in the *dnaA*_T1_-induced cultures and 8.7% of the uninduced exhibited abnormal morphologies (**Fig. 6D**). The parental and empty vector strains, independent of IPTG, did not exhibit such elevated proportions of these cells, indicating that the observed helical abnormalities were due to targeted silencing of the *dnaA* gene. The high level of abnormal cells in the uninduced *dnaA*_T1_ cultures, relative to the controls, can again be explained by leaky expression of the *dnaA* sgRNA encoded in the CRISPRi construct.

**Figure 6.**
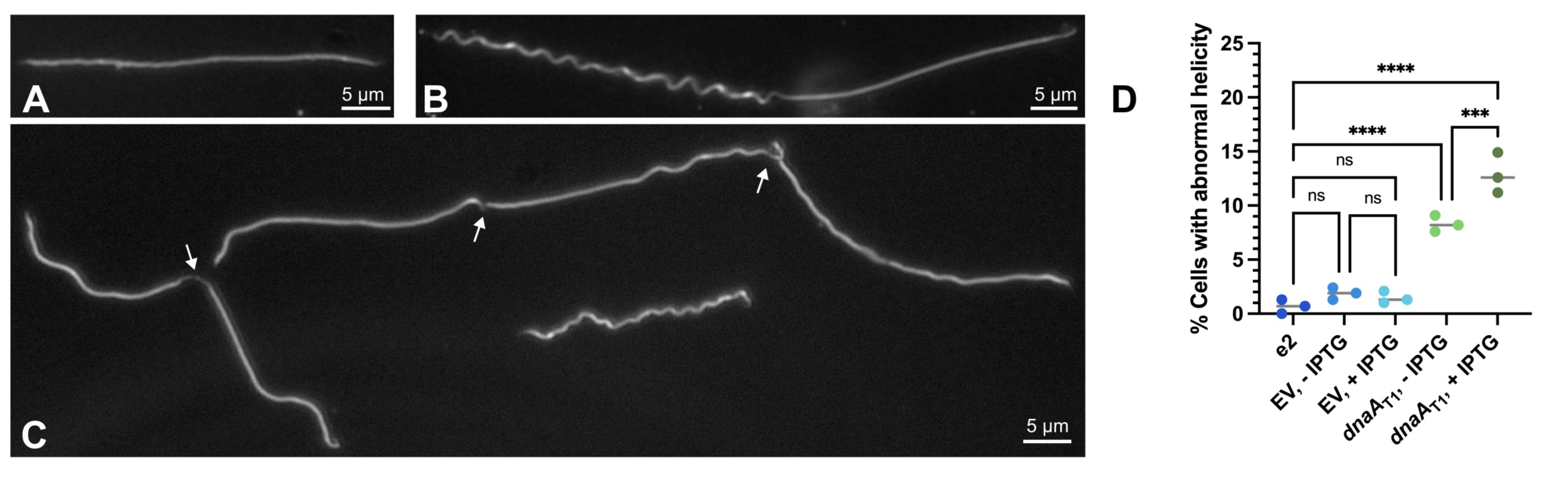
DnaA-depletion affects borrelial helicity. Representative micrographs of observed phenotypes. When *dnaA* was knocked down, spirochetes with no **(A)** or asymmetrical helicity **(B)** were observed. **(C)** Some of the abnormal cells were elongated and showed division defects. **(D)** Spirochetes exhibiting similar defects were observed at a low level in the B31-e2 strain (0.7%, *n* = 1514). The empty vector (EV) control cultures had a similar proportion, independent of the addition of IPTG (uninduced: 2.1%, *n* = 1439; induced: 1.6%, *n* = 1608). The bacteria with the *dnaA*_T1_ CRISPRi plasmid had significantly more cells with perturbed helicity that was dependent on induction (uninduced: 8.7%, *n* = 1437; induced: 13%, *n* = 1472). Spirochetes were counted from three independent cultures. Statistical significance was determined by one-way ANOVA with a Holm-Šídák’s post-hoc test, and multiplicity-adjusted p-values are reported. ***, p ≤ 0.001, ****, p ≤ 0.0001; ns, not significant, p > 0.05.

The characteristic corkscrew appearance of *B. burgdorferi* is due to the 7-11 flagella found in the periplasm (28). The motors that direct the movement of the endoflagella are normally localized to the bacterium’s poles (29). The observed impacts on growth and partitioning suggest coordination between replication and flagellar assembly at the sites of division. This is not unheard of. In *E. coli*, for example, DnaA is known to be involved in flagellar regulation (30, 31).

### DnaA is a global regulator of borrelial gene expression

DnaA is well known to be a transcription factor in many other bacteria, with regulons driving specific phenotypes (9). In *B. burgdorferi*, we previously demonstrated that DnaA controls the expression of the *dnaX-ebfC* operon (10). Considering this and the dramatic impacts of DnaA dysregulation on the Lyme spirochete, we queried whether the myriad phenotypes of the conditional mutants are due to DnaA-dependent transcriptional effects. To address this question, we performed RNA-seq on three strains with different levels of DnaA enrichment: wild-type (e2), DnaA-up (e2 + pACK121), and DnaA-down (e2 + *dnaA*_T1_). All the strains had the plasmids cp26, lp17, lp54, cp32-1, cp32-3, and cp32-4, as assessed by PCR and whole genome sequencing. The bacteria for RNA-seq analysis were grown to mid-exponential phase, induced with 0.5 mM IPTG, and incubated overnight. We chose this strategy to mitigate the chance of escape-mutant development from confounding the results.

For the analyses of the RNA-seq data, comparisons were made between the three groups. We considered a gene to be differentially expressed if the false discovery rate (FDR) was less than or equal to 0.05 and had a log_2_FC (fold change) of ≥ 1 or ≤ −1. With these thresholds, 216 genes were noted as being differentially expressed between the wild-type and DnaA-down cultures (72 down, 144 up; **Fig. 7A**), 84 genes between WT and DnaA-up (40 down, 44 up; **Fig. 7B**), and 259 between DnaA-down and DnaA-up (105 down, 154 up; **Fig. 7C**). Principle component analysis showed that the replicates of each strain clustered together (**Fig. 7D**). Most of the genes that were differentially expressed are encoded on the chromosome: DnaA-down vs. e2: 69.9%; DnaA-up vs. e2: 72.6% (**Fig. 7E**). Of the plasmids, the linear lp54 replicon had the most impacted genes for the DnaA-down vs. e2 comparison (8.3%), while the circular cp32-1 had the most for the DnaA-up vs e2 comparison (7.1%; **Fig. 7E**).

**Figure 7.**
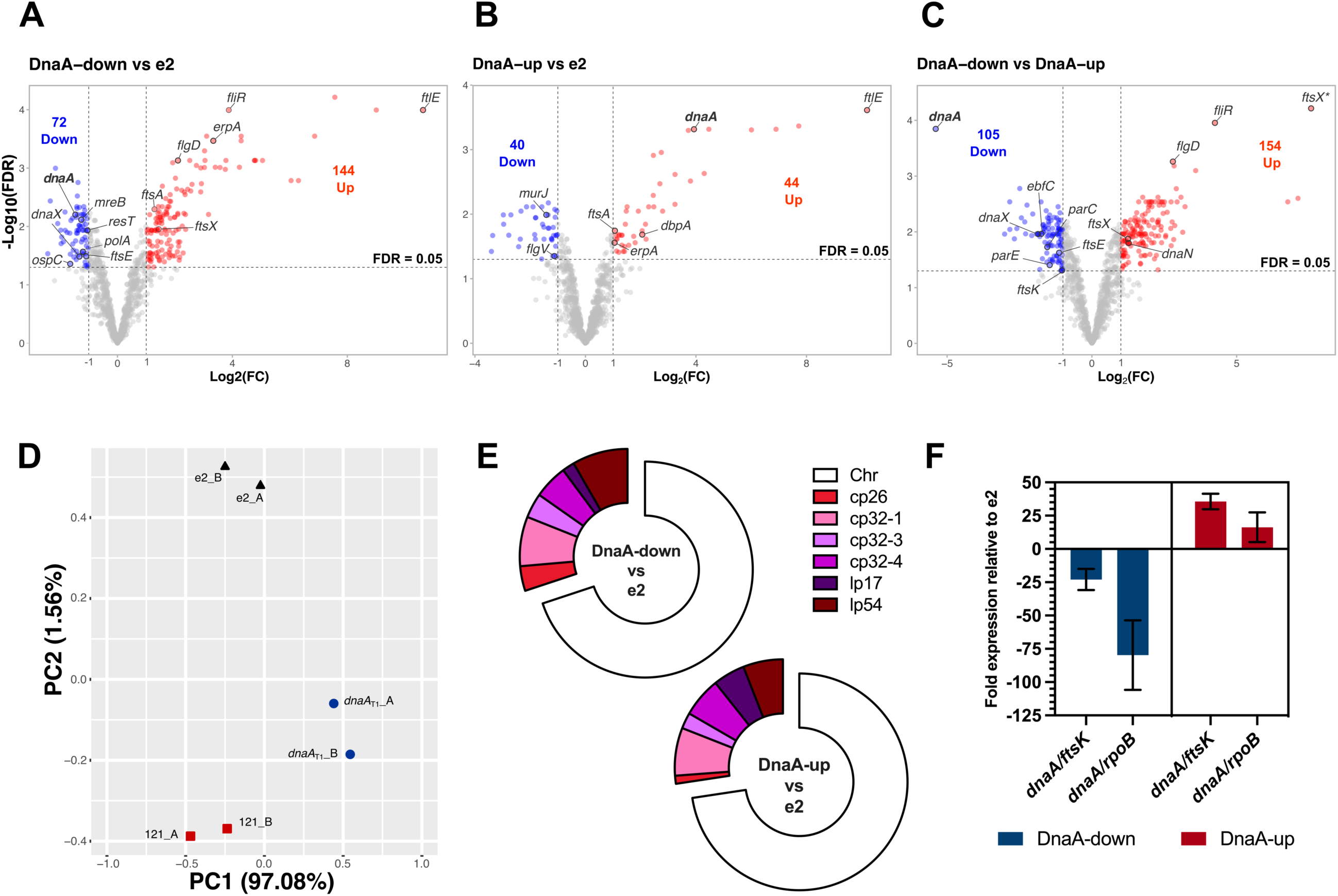
RNA-seq of DnaA-dysregulated *B. burgdorferi*. The RNA from three *B. burgdorferi* strains with different cellular levels of DnaA were sequenced: B31-e2 (parent), CRISPRi *dnaA*_T1_ (DnaA-down), and overexpression pACK121 (DnaA-up). **(A-C)** Volcano plots plotting the fold change (log_2_FC) against each gene’s false discovery rate (-log_10_(FDR)). The threshold for significance was set at a log_2_FC ≥ 1 or ≤ - 1 (≥ 2-fold change) and an FDR of ≤ 0.05. **(D)** PCA plot showing the clustering of the replicate samples that were sequenced. **(E)** Donut charts showing the replicons on which the significantly affected genes are encoded. **(F)** qRT-PCR results of *dnaA* transcripts in the sequenced samples. Cq values were normalized to *ftsK* or *rpoB* (ΔCq) and then to the parental strain (ΔΔCq). Error bars represent the standard deviation (SD).

As expected, the RNA-seq showed that compared to the e2 parent, *dnaA* transcript was about 2.8-fold less abundant in the CRISPRi strain and 15-fold more abundant in the overexpression strain. Typically, we normalize transcript levels to *ftsK*, a gene previously observed to have stable expression across growth phases (32). RNA-seq showed that *ftsK* was impacted by *dnaA* dysregulation (discussed below). Given this, we performed validations normalizing to *rpoB*, which did not change under the tested conditions. Using this approach, *dnaA* transcript was 80-fold less abundant in the knockdown vs. the WT and 16-fold more abundant in the overexpression vs. the WT (**Fig. 7E**). DnaA protein levels were about half as abundant in the knockdown and 13 times more in overexpression than the WT (**Fig. 10B**).

Transcript levels of many genes were most affected when *dnaA* levels were reduced. We initially hypothesized that transcriptomic profiles between the conditional mutants relative to the parental strain would reciprocally mirror each other. Only one gene fit this hypothesis, BB_0413 (2.2-fold increase for *dnaA*_T1_/e2; 2.3-fold decrease for pACK121/e2). Indeed, in many cases, transcript levels for a gene would experience the same degree of change regardless of whether DnaA was increased or decreased. Such instances suggested that some of the observed changes in gene expression were not due to DnaA itself but rather a shared cellular response to the changes in DnaA levels. Despite some of these complications, the RNA-seq data set offered insights that could explain the phenotypes observed in the conditional *dnaA*-mutant strains.

The earliest interrogations into borrelial DnaA focused on its function as a transcription factor (10, 14). Consistent with our prior *in vitro* studies, knocking down *dnaA* significantly decreased transcription of the *dnaX-ebfC* operon (10). This operon codes for the *τ*-subunit of the DNA polymerase III holoenzyme (DnaX) and the nucleoid-associated protein, EbfC. Other impacted replication genes included the DNA polymerase III β-clamp (*dnaN*), DNA polymerase I (*polA*), DNA topoisomerase IV (*parEC*), and telomere resolvase (*resT*). Except for *dnaN*, these genes were downregulated when *dnaA* levels were decreased. The upregulation of *dnaN*, seen when comparing DnaA-Up to DnaA-down conditions, suggests that DnaA represses the expression of this gene. This possibility is unsurprising given that the *oriC* is directly 5’ of the *dnaN* ORF, and we have shown that DnaA binds this region (10).

In the *dnaA*-knockdown *B. burgdorferi*, we observed cells that were considerably elongated. In our evaluation of the borrelial genome, we identified 22 homologs of genes that encode components of either the bacterial elongasome or divisome (**Table 1**). Of these, 39.1% were differentially expressed in the *dnaA*_T1_ CRISPRi strain (**Fig. 8A**). Among these were *mreB*, *mreC*, *mreD,* and *mrdA*, which constitute an apparent operon and encode core components of the bacterial elongasome (**Fig. 8B**). MreB is an actin-like cytoskeleton protein that polymerizes across the bacterial inner membrane to regulate the spatiotemporal peptidoglycan synthesis of growing cells (33–35). Knockdown of MreB in *B. burgdorferi* results in bulging and cell widening at the division sites (19). MreC and MreD are membrane-embedded proteins that interact and regulate the peptidoglycan crosslinking activity of PBP2 (penicillin-binding protein 2; MrdA) (36). The peptidoglycan polymerase RodA and PBP1a interact with PBP2, with the former stimulating its activity. Except for *mreD*, the genes in the apparent *mre* operon were downregulated in *dnaA*-deficient *B. burgdorferi*. A DNA sequence that is the same as those found in the *oriC*, which may constitute the borrelial DnaA-box, is located upstream of the *mreB* ORF, suggesting DnaA may directly regulate the expression of these genes. The decreased transcript levels for *mreB*, *mreC*, and *mrdA* would suggest decreased elongation, which is the opposite of what we observed (**Fig. 3**). This apparent contradiction suggests that filamentation may be due to defects in cell division.

**Figure 8.**
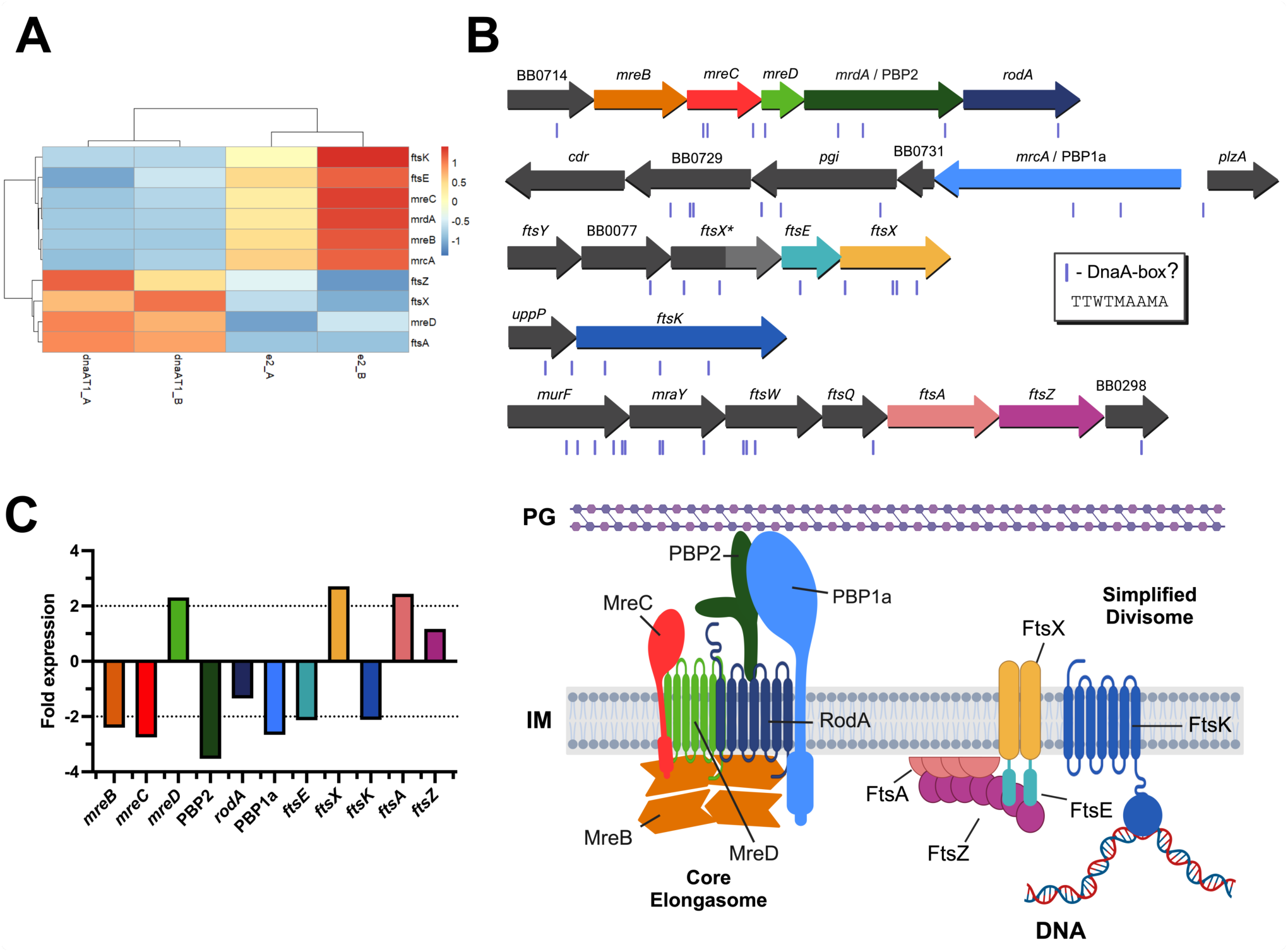
Impacts of DnaA on *B. burgdorferi* elongation and division genes. **(A)** Heat map of the division and elongation genes that were significantly affected when DnaA was knocked down. **(B)** Top: Locus maps of the select elongation and division genes to scale. The sites of potential DnaA-boxes are denoted as purple boxes below the genes. Bottom: Simplified schema of the hypothesized borrelial elongasome and divisome containing the highlighted genes of interest. Informed by the work of Liu et al. and Hu et al. (36, 69) **(C)** Bar graph showing the fold change detected by RNA-seq of the elongation and division genes. The dashed lines indicate the 2-fold threshold for meaningful gene expression changes.

**Table 1.** Homologous elongation and division genes of *B. burgdorferi*.

| Protein | Locus Tag | Elongasome | Divisome | RNA-seq Differential Expression |  |  |
| --- | --- | --- | --- | --- | --- | --- |
|  |  |  |  | DnaA-down<br>vs e2 | DnaA-up<br>vs e2 | DnaA-down<br>vs DnaA-up |
| FtsE | BB_0080 | - | + | Down | NA | Down |
| FtsX | BB_0081 | - | + | Up | NA | Up |
| FtsI/PBP3 | BB_0136 | + | + | NA | NA | NA |
| MurE | BB_0201 | + | + | NA | NA | NA |
| FtsK | BB_0257 | - | + | Down | NA | Down |
| FtsZ | BB_0299 | - | + | NA | NA | NA |
| FtsA | BB_0300 | - | + | Up | Up | NA |
| FtsQ/DivIB | BB_0301 | - | + | NA | NA | NA |
| FtsW | BB_0302 | - | + | NA | NA | NA |
| MraY | BB_0303 | + | + | NA | NA | NA |
| MurF | BB_0304 | + | + | NA | NA | NA |
| MurD | BB_0585 | + | + | NA | NA | NA |
| MurB | BB_0598 | + | + | NA | NA | NA |
| Ami | BB_0666 | - | + | NA | NA | NA |
| MreB | BB_0715 | + | - | Down | NA | Down |
| MreC | BB_0716 | + | - | Down | NA | Down |
| MreD | BB_0717 | + | - | Up | NA | Up |
| PBP2 | BB_0718 | + | - | Down | NA | Down |
| RodA | BB_0719 | + | - | NA | NA | NA |
| PBP1a | BB_0732 | + | - | Down | NA | Down |
| MurG | BB_0767 | + | + | NA | NA | NA |
| MurJ | BB_0810 | + | + | NA | Down | Up |
| MurC | BB_0817 | + | + | NA | NA | NA |

\* Plus sign (+) indicates the protein is part of the complex, minus sign (-) indicates it is not
\* NA = Not affected

The transcripts of four important divisome proteins were impacted when *dnaA* was knocked down: FtsA, FtsK, FtsE, and FtsX (**Fig. 8A** and **B**). FtsA is the membrane anchor protein for the septation marker protein FtsZ (37, 38), and its transcript increased about 2-fold in the conditional *dnaA* mutant. Surprisingly, DnaA-overexpressed *B. burgdorferi* also experienced a 2-fold increase in *ftsA*. Transcripts for FtsK, the DNA translocase, declined 2-fold. FtsE and FtsX, encoded next to each other, had their transcripts decreased 2-fold and increased almost 3-fold, respectively. Interestingly, a second allele of *ftsX* (*ftsX\**), which is truncated, is located directly upstream of *ftsEX,* and transcript levels of this gene were substantially affected, with a 189-fold increase. Like the *mre* locus, DnaA could regulate these genes directly as potential DnaA-boxes are adjacent to these loci.

In the conditional *dnaA*-knockdown *B. burgdorferi,* we observed spirochetes that had lost their helical structure and hypothesized these cells had disrupted flagella. The *B. burgdorferi* genome encodes nearly three dozen flagellar genes (**Fig. 9A**). When *dnaA* was knocked down, six of these genes were significantly impacted: *fliQ*, *fliR*, *flgD*, *flaA*, *flgV*, and *flgA* (**Fig. 9B**). All these genes except for *flaA* and *flgA* are encoded in the *flgB* superoperon. FliQ and FliR are components of the flagellar export apparatus (39). FlgD is a scaffolding protein that is required for flagellar hook formation (40, 41). FlaA is the minor borrelial flagellar filament protein (42, 43). FlgV is a protein first characterized as a homolog of the RNA chaperone Hfq, but recent work has demonstrated the protein localizes with flagellar motors and modulates their assembly (44, 45). FlgA is a chaperone involved in the formation of the P-ring of the flagellar motor (46). The transcripts for *fliR* and *flgD* increased the most of these genes, about 15 and 4-fold, respectively, when DnaA levels were reduced. The changes in the expression of these genes could alter flagellar assembly homeostasis and explain the observed abnormalities in helicity in the *dnaA*-deficient spirochetes. It is also possible that defects in septation disrupted the localization and assembly of flagella.

**Figure 9.**
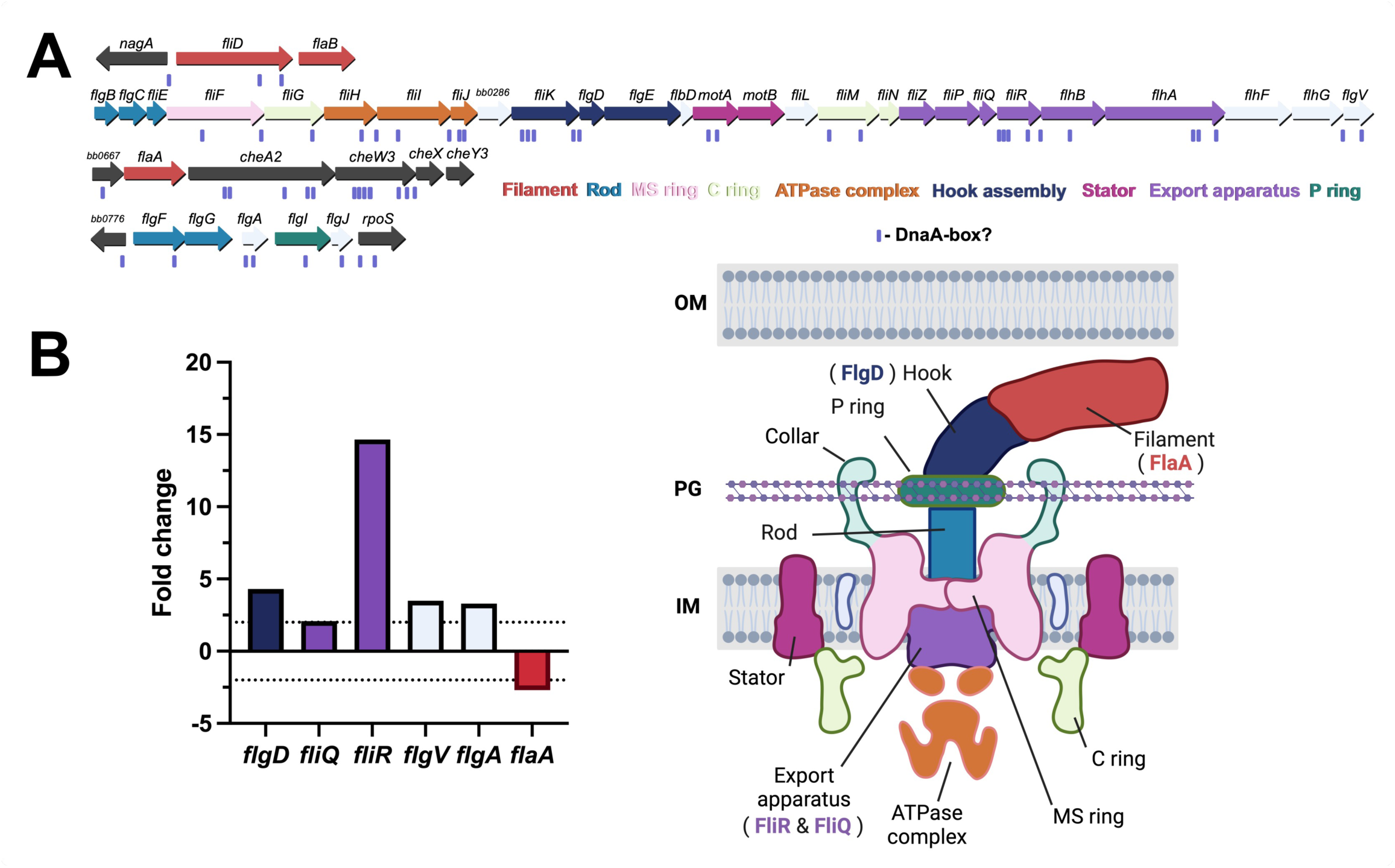
Impacts of DnaA on *B. burgdorferi* flagellar genes. **(A)** The borrelial flagellar motor is a complex molecular machine (bottom right) that is comprised of nearly three dozen proteins encoded in four separate loci on the chromosome (top). The schematic of the *B. burgdorferi* flagellar motor was informed from data by Zhao et al., Qin et al., and Chang et al. (98–100). **(B)** RNA-seq showed the transcripts for five flagellar genes increased when DnaA was knocked down. Two of these genes, FliQ and FliR (purple), are constituents of the flagellar export apparatus. The dashed lines indicate the 2-fold threshold for meaningful gene expression changes.

In addition to affecting genes for essential cellular processes, DnaA significantly impacted the expression of outer surface proteins that are involved in vertebrate infection and virulence, such as ErpA, decorin binding protein DbpA, and OspC (**Fig. 7A-C** and **Fig. 10E-F**). The Erps are multifunctional adhesins that bind vertebrate host factors such as laminin, plasminogen, and complement proteins (47–52). Transcripts for *erpA* increased in the *dnaA* knockdown and overexpression conditions (**Fig. 10E-F**). Interestingly, this increase in transcript didn’t correspond to increased ErpA protein (**Fig. 10B-D**).

**Figure. 10.**
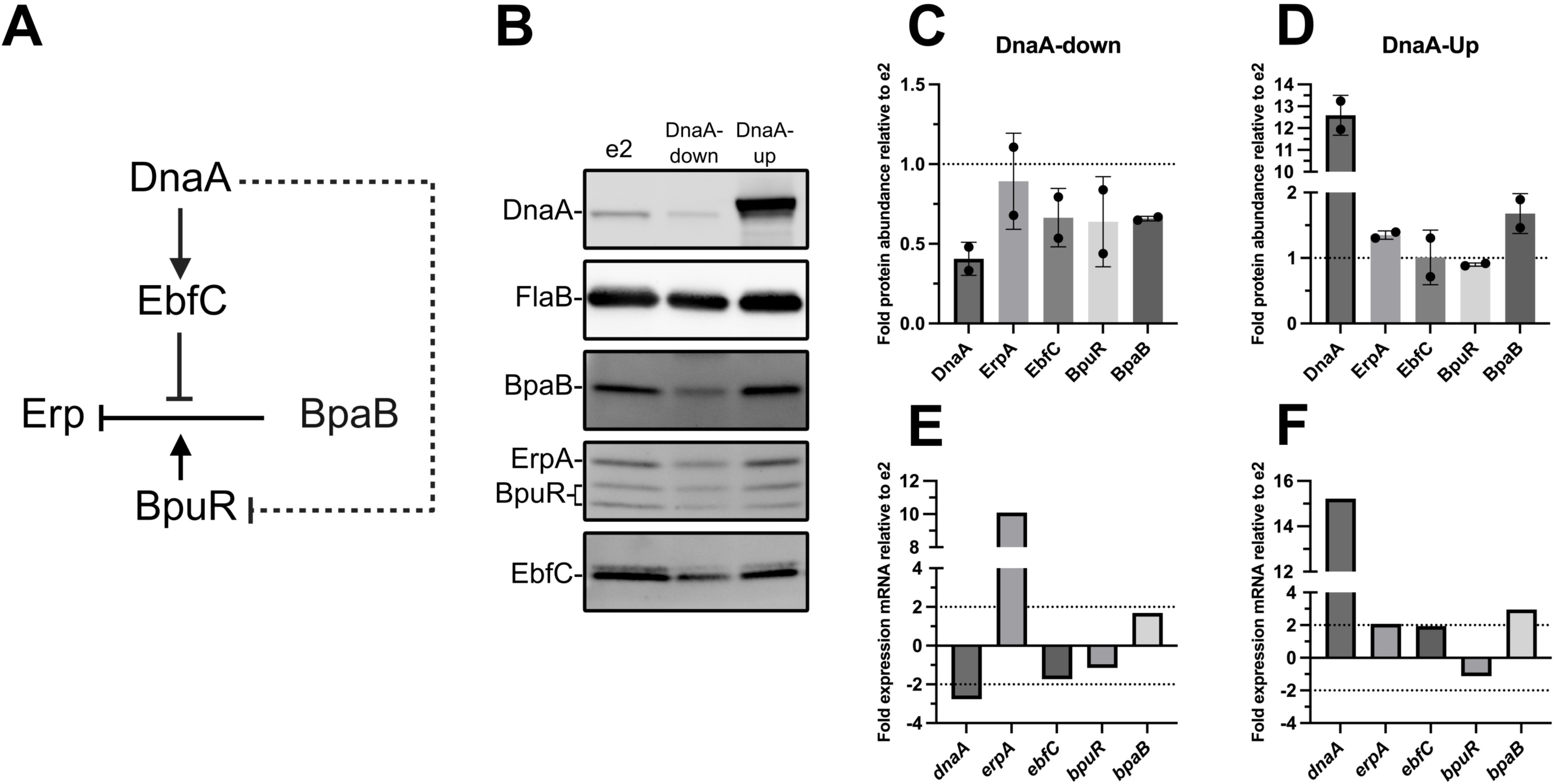
Impacts of DnaA on the Erp regulatory network. **(A)** Solid black lines indicate established interactions. Dashed lines indicate hypothesized interactions. Activation is denoted by lines with arrowheads, and inhibition by lines without arrowheads. Note that co-repressive or anti-repressive activities, i.e., BpuR and EbfC, respectively, are indicated by arrows directed at lines. **(B)** Representative immunoblots from samples of *B. burgdorferi* that were analyzed by RNA-seq. Quantitation of target proteins when DnaA was knocked down **(C)** or overexpressed **(D)**. Band intensities for each replicate immunoblot were normalized to FlaB and then the parental e2 strain. **(C-D)** The dotted lines at 1 represent where the protein abundance is the same as e2. Bar graphs showing the fold change detected by RNA-seq in the DnaA-down/e2 **(E)** and DnaA-up comparisons **(F)**. **(E-F)** The dotted lines indicate the 2-fold threshold for meaningful gene expression changes.

The *erp* genes are encoded on the cp32 plasmids and have conserved operators that allow uniform regulation (53). The Erps are turned on when the tick begins to feed, a time of rapid borrelial replication, and remain on during vertebrate infection (54). Three proteins bind the *erp* operators and regulate their expression: BpaB (<u>B</u>orrelial cp32 P<u>arB</u> analog), BpuR, and EbfC. BpaB represses *erp* transcription along with the co-repressor BpuR (**Fig. 10A**) (55–57). EbfC is the antirepressor that antagonizes BpaB to allow for *erp* transcription (56). DnaA was previously hypothesized to regulate *erp* transcription by activating EbfC and repressing BpuR (10, 13, 14). EbfC transcript and protein levels decreased when *dnaA* was knocked down, consistent with that model (**Fig. 10B** and **E**). BpuR transcript, however, did not significantly change, but its protein levels appeared to decline with a decrease in cellular DnaA concentrations (**Fig. 10B** and **E**). BpuR alone does not repress *erp* expression, so this cannot account for the increase in ErpA transcript. Thus, we looked at the protein levels of the *erp* repressor. Despite the lack of a change in the transcript, immunoblots showed a decrease in BpaB protein levels in *dnaA* CRISPRi *B. burgdorferi*, which can explain the increase in *erpA* transcript (**Fig. 10B, C**, and **E**). BpaB levels were similar to those of the parental e2 strain during *dnaA* overexpression, and ErpA transcript and protein were unaffected, likely due to normal levels of the antirepressor EbfC. Overall, these results demonstrate that DnaA not only affects the expression of genes involved in basic cellular processes but also those involved in maintaining the Lyme spirochete’s infection cycle.

## DISCUSSION

In this study, we sought to understand the roles of DnaA in controlling the physiology of *B. burgdorferi*. To do this, we utilized inducible CRISPRi and overexpression vectors. Overexpressing DnaA was toxic to the bacteria and was consistently overcome by mutations in the inducible *dnaA*. This, along with the difference in DnaA levels from overexpression vs. knockdown, 13-fold DnaA increase vs. 2-fold decrease, likely explains some of the results we encountered. Nevertheless, increasing DnaA levels yielded significantly different phenotypes, such as spirochete size. CRISPRi was the more reliable and effective means of producing conditional *dnaA* mutants in *B. burgdorferi*. This approach targeting *dnaA* has been successfully described in *E. coli*, *Lactobacillus plantarum,* and *Streptococcus pyogenes* (58–60). An attempt was made in *Pseudomonas putida,* but clones couldn’t be recovered (61).

We found that lowering the cellular levels of DnaA profoundly reduced cell division, consistent with the protein’s function as the chromosomal replication initiator. The reduction of DnaA also coincided with decreased expression of many essential replication genes. It is known that one of these genes, *dnaX*, is directly regulated by DnaA (10). This suggests that DnaA may directly regulate those other genes, or *B. burgdorferi* may have mechanisms to sense replication initiation and coordinate gene expression accordingly.

DnaA-deficient spirochetes also increased in length. The bacterial cell cycle is typically divided into three phases: B, C, and D (62). The B and D phases correspond to the <u>b</u>irth and <u>d</u>ivision of the bacteria, respectively. The C phase corresponds to everything in between, namely, <u>c</u>hromosomal replication, along with elongation/growth. Replication is initiated by DnaA and then carried out by the replisome. Elongation is facilitated by the elongasome, which synthesizes peptidoglycan to allow for cell growth. The DnaA-depleted spirochetes, although considerably longer than the parental strain, had overall decreased expression of elongasome genes, the *mre* locus in particular. This suggested a division issue, but we cannot rule out the possibility that DnaA is required for cross-talk between replication and cell growth. Indeed, it is likely that replication, elongation, and division are all interconnected. Lyme and relapsing *Borreliae* elongate at three distinct zones along their length: 1/4, 1/2, and 3/4 (63). This localization pattern at the mid-cell suggests an interplay between elongation and division machinery. How or if DnaA directly plays into this delicate balance remains to be assessed.

Many bacterial cell division proteins are denoted “Fts” for their filamentous phenotypes caused by temperature-<u>s</u>ensitive mutations (64). Knocking down DnaA affected the expression of *ftsA*, *ftsEX*, and *ftsK.* The dysregulation of these genes potentially explains both the elongated phenotype and the irregular spacing of the ParB-*oriC* puncta.

FtsA localizes to the inner leaflet of the inner membrane and binds FtsZ. Increased FtsA, seen when DnaA was knocked down, could decrease Z-ring assembly and, thus, cytokinesis. In *E. coli*, overexpression of FtsA causes cells to filament (65). Overexpressed DnaA spirochetes had elevated *ftsA* transcript, but no changes were observed in *ftsEX*. The FtsEX complex regulates the activity of peptidoglycan hydrolases and FtsA (66–69). Disruption of FtsE and FtsX levels could, therefore, prevent the breakdown of peptidoglycan and Z-ring formation at the site of division. The combined perturbation of *ftsA* and *ftsEX* expression in the *dnaA* CRISPRi strain potentially explains the observed elongated phenotype.

In addition to changes in spirochete length in the DnaA-depleted *B. burgdorferi*, we observed abnormal spacing of ParB-*oriC* puncta. These bacteria had decreased transcripts of the membrane-embedded DNA translocase FtsK, a core component of the divisome (70–72). FtsK, through its C-terminal αβγ-domain, resolves chromosome dimers and translocates DNA from the septum to allow for cytokinesis (73–76). A decrease in FtsK could thus result in the failure to separate sister chromosomes, leading to cells with abnormal ploidy. This could also explain why some *oriC* sites were located close together in the *dnaA* knockdown strain (**Fig. 5**). FtsK also recruits downstream divisome proteins through its N-terminal transmembrane domain (71, 77–79). Therefore, reducing FtsK may also limit the recruitment of divisome proteins to the septa.

Motility is perhaps the most important “virulence” factor that *B. burgdorferi* employs to infect and colonize its vertebrate hosts. By knocking down DnaA in the Lyme spirochete, we observed a substantial impact on the bacteria’s helicity. This corkscrew morphology is due to periplasmic flagella (28). Key genes, such as FlgD and FliR, were overexpressed when levels of DnaA were lowered. FliR is a component of the flagellar motor’s export apparatus, which transports flagellar filament substrates from the cytoplasm to the periplasm (39). Disrupting the balance of this subunit would very likely impair the overall function and structure of the flagellar motor and filament. By increasing FliR, it is possible that the export of the flagellar cargo to the periplasmic space would be affected, which in turn would prevent flagella formation. The flagellar motors are normally localized to the poles and developing septa of *B. burgdorferi*. This suggests that replication, elongation, or division processes may coordinate the formation of flagellar motors at the dividing mid-cell. In *E. coli*, DnaA regulates genes involved in flagellar assembly (30, 31, 80). Whether DnaA in *B. burgdorferi* acts through a similar mechanism is unknown.

In addition to the many morphological changes incurred by dysregulation of DnaA, there were also significant impacts on the expression of vertebrate infection-related outer surface proteins. Many of these antigenic proteins, such as OspC, DbpA, and the Erps are turned on during the tick blood meal, a period defined by rapid replication. Relative to these genes, levels of transcript for the alternative sigma factor RpoS were not significantly changed by manipulation of DnaA levels. We previously demonstrated that DnaA regulates the *erp* antirepressor EbfC and may regulate the *erp* co-repressor BpuR (10, 14). With our CRISPRi approach, we were able to validate our prior data on *dnaX-ebfC* regulation. While we could not detect any substantial differences in BpuR with DnaA manipulation, this might have been a consequence of assessing mid-exponential spirochetes grown at 34 °C; BpuR expression is highest when grown at 23 °C (14, 57). When DnaA levels were reduced, expression of ErpA increased. We suggest this could be due to decreased levels of *erp* repressor BpaB. While this was unexpected, it is logical that a partitioning protein be decreased when *dnaA* transcription is knocked down. This suggests that these extrachromosomal DNAs can sense changes in chromosomal replication. How this is mediated remains to be investigated.

We observed gene expression level changes in a vast number of regulatory networks. This led us to examine if *dnaA* knockdown or overexpression impacted any known regulatory factors. Among the known networks, we observed modest changes in gene expression levels of the following regulators: Hk1, Rrp1, and SpoVG. These changes did not meet the log fold change threshold but were significant (FDR ≤ 0.05) and had a log fold change of approximately 1.5. When comparing the *dnaA*_T1_ knockdown strain to the parental e2 strain, these genes were downregulated by a fold change of 1.52, 1.61, and 1.53, respectively. While not meeting our set threshold, it is possible the combined modest expression changes in these regulators considerably impacted the expression levels of the genes they modulate. For example, the nucleic acid binding protein SpoVG, the response regulator Hk1, and diguanylate cyclase Rrp1 are all known to regulate the glycerol metabolism (*glpFKD*) operon, which is essential for *B. burgdorferi* colonization and persistence in ticks (81–85). The downregulation of the *glpFKD* operon (∼2.6 to 4-fold) is likely attributable to the decreased expression of *spoVG*, *hk1*, and *rrp1*. These observations link DnaA and/or DNA replication to crucial borrelial networks, including the c-di-GMP regulatory network (Hk1/Rrp1) critical to completing the *B. burgdorferi* enzootic life cycle.

Another regulatory network impacted was the Hk2/Rrp2 pathway, whose expression levels were increased by 8.2 and 1.5, respectively. The Hk2/Rrp2 two-component system is a known activator of the RpoN/RpoS alternative sigma factor cascade, which regulates essential virulence factors like OspC (86–88). Deletion of *rrp2* is lethal, and investigations into *rrp2* regulation have relied on conditional mutants (18). Although neither *rpoN* nor *rpoS* levels changed significantly, we note that *rpoS* transcription can be activated by the housekeeping sigma factor RpoD in addition to RpoN (89). This further provides strong evidence that the Hk2/Rrp2 pathway has impacts outside of *rpoS* regulation potentially mediated by proper *dnaA* levels and/or associated with DNA replication. In addition, this data supports our past data showing that *ospC* regulation, through the repressor Gac, can occur independently of RpoS regulation (90). Additional data from our lab indicates that DnaA is a positive activator of OspC (unpublished results). This supports the RNA-seq data in this study, showing that knockdown of DnaA levels reduces *ospC* transcript.

Organisms have evolved complex and varied molecular circuits to regulate cellular homeostasis in ever-changing environments. Bacterial pathogens, especially, have developed such networks in their tussle against host defenses. During the tick blood meal, *B. burgdorferi* must take advantage of the nutrients to propagate its numbers, reprogram its transcriptome, and disseminate to the vertebrate. Failure to do so means death, as *B. burgdorferi* cannot be transmitted vertically to tick offspring (91). In *B. burgdorferi,* DnaA, the master regulator of replication initiation, appears to form a complex regulatory network to coordinate these essential processes (**Fig. 11**). Future studies need to be conducted to determine which of the DnaA-dysregulated effects described here are due to DnaA itself or a consequence of a lack of initiation of chromosomal replication.

**Figure 11.**
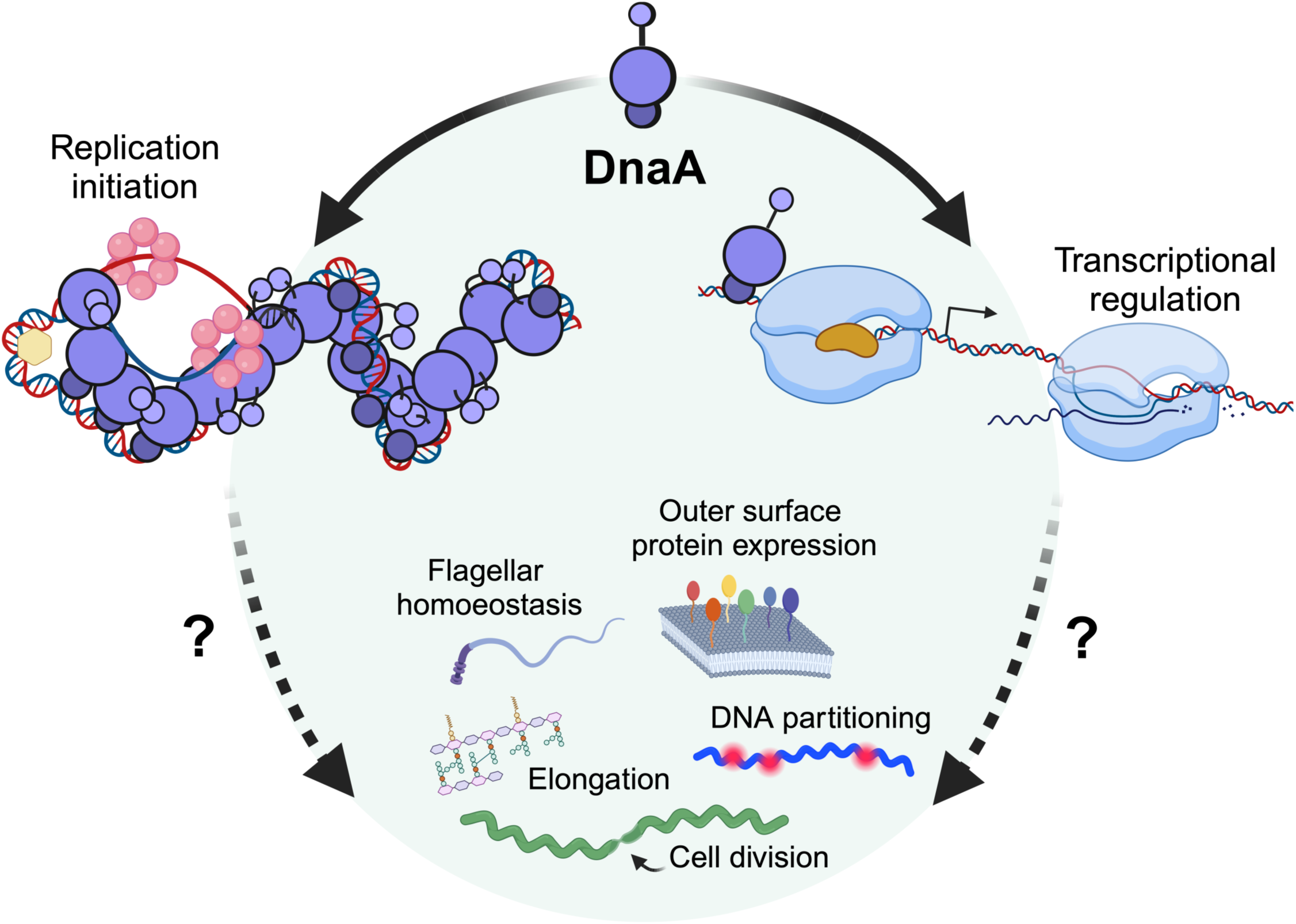
Roles and impacts of DnaA dysregulation on borrelial physiology. The DnaA of *B. burgdorferi* functions as the chromosome replication initiator and a transcription factor. Altering levels of this essential protein modulate aspects of DNA replication and bacterial physiology. The data presented here point to crosstalk between the systems controlling replication and bacterial morphology. Future studies will need to be conducted to ascertain the direct roles DnaA plays in maintaining this delicate balance.

## METHODS AND MATERIALS

### Bacterial strains, plasmids, and genetic manipulations

Studies were conducted using the readily transformable *B. burgdorferi* clonal strain B31-e2 (92). Spirochetes were grown in liquid Barbour-Stoener-Kelly II (BSK-II) medium supplemented with 6% rabbit serum (v/v) at 35 °C. For all experiments, *B. burgdorferi* was diluted from mid-exponential phase (3-5 x 10^7^ cells/mL) into fresh BSK-II medium. Cultures were only used if they were at least two passages away and no more than three from the initial −80°C stock. Borrelial culture densities were enumerated using a Petroff-Hauser counting chamber and dark-field microscopy.

The *dnaA* overexpression (pACK121) and CRISPRi plasmids (pACK136, *dnaA*_T1_; pACK138, *dnaA*_NT1_) were generated by GenScript. The pACK121 plasmid is a derivative of pJSB268 (17). The plasmid was designed to replace the *luc* gene with the *B. burgdorferi dnaA* gene containing an N-terminal 3xFLAG epitope. The pACK136 (*dnaA*_T1_) and pACK137 (*dnaA*_NT1_) plasmids are derivatives of pJJW101 (20). These plasmids were created by inserting the desired sgRNA sequence between the BsaI sequences. The sgRNA targets used in this study were chosen using CRISPy-web (https://crispy.secondarymetabolites.org) and B31 chromosome contig (NC_001318). Whole-plasmid sequencing was performed by Plasmidsaurus using Oxford Nanopore Technology with custom analysis and annotation.

### Growth curve analysis

Growth curves for each strain were generated from three independent experiments. Cultures were seeded at the same initial density (1 or 5 x 10^5^ cells/mL) and enumerated over seven days. Generation times were calculated in Microsoft Excel by fitting an exponential curve to the data points denoting the beginning and end of logarithmic growth. Statistically significant differences were determined by one-way ANOVA.

### Immunoblot analyses

Polyclonal DnaA antibodies were generated by Thermo Scientific in rabbits using recombinant GST-DnaA generated in-house (10). The serum from the final bleed was purified by affinity purification using MBP-DnaA on an amylose column. Murine monoclonal anti-FlaB antibodies were used to assess the even loading of SDS-PAGE gels (93). Other primary antibodies were generated previously (56, 57, 94). Cells for immunoblot were pelleted and washed three times with 1x PBS, pH 7.4. Approximately 10^7^ spirochetes were loaded per lane for SDS-PAGE. Goat anti-Rabbit Alexa Fluor 488 (1:10, 000, Thermo Scientific) and Goat anti-Mouse IRDye800 (1:5,000, Licor) were used as secondary antibodies. Densitometric analyses were performed using ImageLab software (BioRad). The intensities of the bands of interest were normalized to the corresponding loading control.

### Quantitative reverse transcription-PCR (qRT-PCR)

Mid-exponential phase (3-5 x 10^7^ cells/mL) bacteria were washed three times with PBS (pH 7.4) before RNA extraction using Qiagen Mini RNA kits. Genomic DNA was cleared by on-column DNase treatment. RNA was reverse transcribed using iScript cDNA synthesis kits (Bio-Rad). Quantitative PCR was conducted using iTaq Universal SYBR Green Supermix (Bio-Rad) and a Bio-Rad CFX96 Touch Real-time PCR thermocycler. The primer sets used for this study were designed using the IDT Primer Quest Tool (https://www.idtdna.com/PrimerQuest/Home/Index). Cq values were normalized to *ftsK* or *rpoB* (ΔCq) and then to the parental strain or uninduced control (ΔΔCq). Fold expression was determined using the function 2^-ΔΔCq^. If ΔΔCq > 0, then the function −2^ΔΔCq^ was used.

### Cell length analysis

Wet mounts of live bacteria were visualized by dark-field with an Olympus Bx51 microscope at 400x total magnification. Micrographs were taken using a C-mounted Accu-scope Excelis HD camera. Cell lengths were determined using Captavision+ software. A conversion factor of 13.80 pixels/µm was used for all measurements. For each replicate and time point, approximately 50 spirochetes were measured.

### Fluorescence microscopy and image analysis

Spirochetes were pelleted by centrifugation, washed twice with PBS (pH 7.4), and diluted to a final concentration of 1 x 10^5^ cells/mL. Ten microliters of cells were mounted on a glass slide. Cells were visualized using an Olympus Bx51 microscope with a Cool LED p-E300 illumination system. Micrographs of dark-field and red fluorescence were captured for each field of view using a C-mounted Accu-scope Excelis HD camera. Micrographs were merged in ImageJ, and the background was inverted using EZreverse (https://amsterdamstudygroup.shinyapps.io/ezreverse/) with the HSL color space option (95).

### Whole Genome Sequencing (WGS) and analysis

The parental *B. burgdorferi* B31-e2, *dnaA*_T1_ CRISPRi, and *dnaA*-overexpression strains were sequenced to verify plasmid content and determine if mutations occurred in the *dnaA* sequence of the generated strains. Briefly, the strains were grown in duplicate to mid-exponential phase (3-5 x 10^7^ cells/mL). Two mLs of cells were pelleted and washed twice with PBS (pH 7.4) and frozen at −80 °C. DNA extraction and whole genome sequencing were performed by SeqCenter. Briefly, the cells were sent to the SeqCenter facility. DNA was extracted from the pellets using the ZymoBIOMICS^TM^ DNA Miniprep Kit according to the manufacturer’s protocol. DNA concentrations were determined by Qubit.

Sequencing libraries were prepared via the tagmentation-based and PCR-based Illumina DNA Prep kit and custom IDT 10 bp unique dual indices (UDI) with a target insert size of 320 bp. The sequencing was performed either in one or more multiplexed flow-cell runs on an Illumina NovaSeq 6000 sequencer resulting in 2×151 bp paired-end reads. Quality control steps including demultiplexing and adapter trimming were performed with bcl-convert (v4.1.5). Reads were aligned to the *B. burgdorferi* B31 genome assembly (GCF_000008685.2_ASM868v2). Variant calling was performed using BreSeq (v. 0.38.1) under default settings.

### RNA sequencing (RNA-seq) and analysis

The same cultures of the parental *B. burgdorferi* B31-e2, *dnaA*_T1_ CRISPRi, and *dnaA*-overexpression strains used for WGS were used for RNA-seq. All strains were grown in duplicate to mid-exponential growth phase (3-5 x 10^7^ cells/mL) and induced with 0.5 mM IPTG overnight at 35 °C. Nine mLs of the cells were pelleted and washed two times with PBS (pH 7.4) and frozen at −80 °C. The cells were sent to SeqCenter for RNA extraction and sequencing using their intermediate RNA analysis with replicates (Prokaryotic) service. Briefly, their method consisted of quality control and adapter trimming by bclconvert (version 4.1.5) and mapping reads using HISAT2 (version 2.2.0) to the RefSeq version of the *B. burgdorferi* B31 genome assembly (GCF_000008685.2_ASM868v2). The read counts were uploaded into R (version 4.0.2) and normalized using the edgeR Trimmed Mean of M values (TMM) algorithm (1.14.5). Differential gene expression analysis was performed with edgeR using the normalized TMM values. Only plasmids that were confirmed to be present were included in the analyses. We further filtered the data set by setting the thresholds for differential expression at log_2_FC ≥ 1 or ≤ −1 and an FDR ≤ 0.05. Differentially expressed genes (DEGs) were visualized via volcano plots made using VolcaNoseR (96). DEGs belonging to the elongasome and divisome were visualized by heatmap. The heatmap was generated in R (v 4.4.1) using the pheatmap package, with the log2 transformed normalized counts per million (CPMs) (97). The clustering method used for the heatmap was the default method of complete linkage, while the distance measure used was correlation.

To identify flagellar, cell division, and elongation homologs, the genome was directly searched for annotations, or the sequences of characterized proteins from other bacteria were aligned by BLAST against the *B. burgdorferi* genome.

## Acknowledgments

This research was funded by NIH grant R21 AI147139. We thank Dr. Wolfram Zückert and Bryan Murphy for their help designing and troubleshooting the CRISPRi shuttle vectors and Dr. Jon Blevins for providing the pJSB268 shuttle vector. BioRender was used to make the schematics in Figures 1, 2, 8, 9, and 10.

---

**Table 2.**
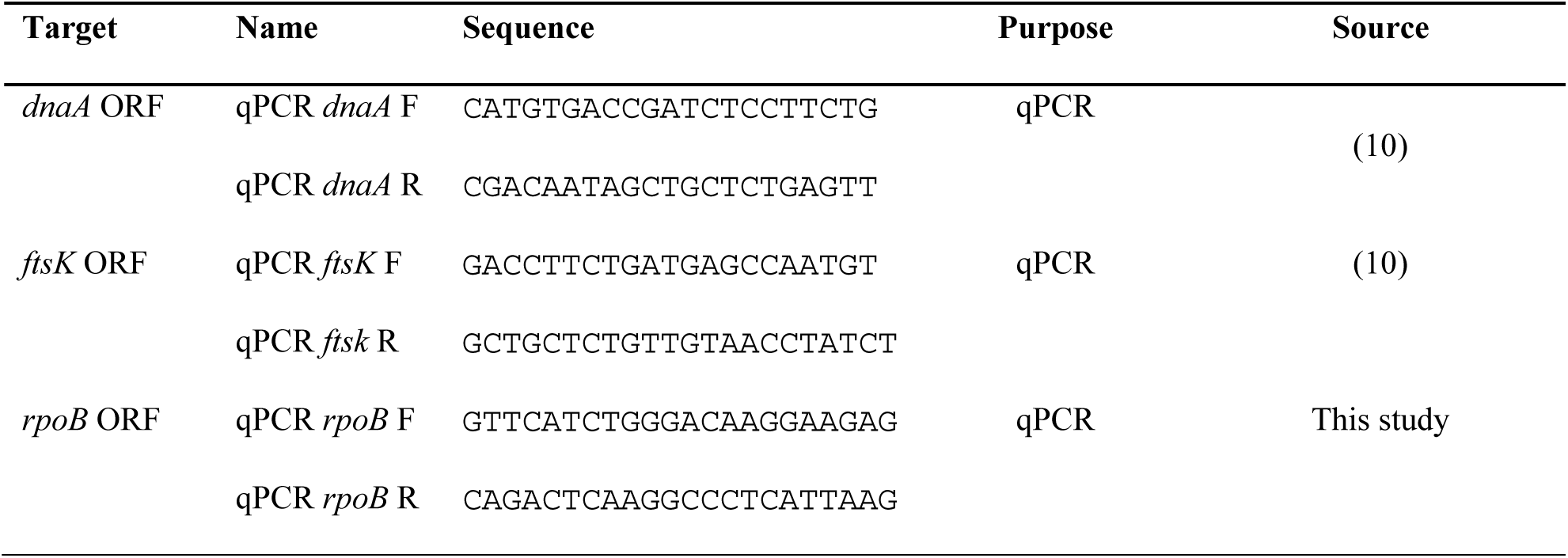
Oligonucleotides used in this study.

